# The diversity of disease-associated natural human antibody specificities declines dramatically in Alzheimer’s disease

**DOI:** 10.64898/2026.07.21.739870

**Authors:** J.M. Reyes-Ruiz, J. Zhang, J. Netanel, Z. Yu, C. G. Glabe

## Abstract

Alzheimer’s disease (AD) is the leading cause of dementia and administration of monoclonal antibodies against Aβ amyloid are associated with disease modifying effects leading to FDA approval for some antibodies. Naturally-occurring human antibodies (NAbs) are believed to be the front line of defense against a variety of diseases, so we investigated the specificity of NAbs in AD and cognitively normal (NC) individuals using epitomic profiling (EP). We found that the diversity of antibody-specific peptide epitope segments (ES) declines dramatically (26-fold) in the AD compared to NC populations. Many of these differentially expressed ES (DEES) map to amyloid sequences that accumulate in AD, such as Aβ, tau, α-synuclein, TDP43 and TMEM106b. NAbs that target amino terminal Aβ epitopes are elevated in AD (residues 1-6 and 18-21) and NAbs that target different Aβ epitopes are elevated in NC (residues 5-10 and 35-41). DEES that map to 2N4R tau are highly cross correlated in the population implying that that the antibodies may be polyspecific or that a set of antibodies are coordinately regulated. We used weighted gene correlation network analysis (WGCNA) to identify co-expressed DEES and UMAP to embed DEES in both AD and NC populations that bind to the same antibody or co-expressed set of antibodies. An average of approximately 25 DEES cluster in each WGCNA group. A major WGCNA cluster associated with AD contains 3 groups of overlapping DEES that assemble into longer linear epitopes. One of these 3 groups contain sequences that map to the amino terminus of Aβ. DEES elevated in AD may be useful blood biomarkers for AD. The DEES clusters elevated in NC may be associated with protection against misfolded protein amyloid seeding and propagation and maintaining the resilience of neuronal populations.

## Introduction

Alzheimer’s disease (AD) is a neurodegenerative disorder characterized by accumulation of amyloids in the brain and a loss of normal neurological functional abilities leading to memory and cognitive impairments, and progressive behavioral changes. AD is the leading cause of dementia with more than 5.8 million people diagnosed in the US and this number is expected to reach over 13 million by 2050 (1). AD pathology includes the accumulation of amyloid fibrils formed from two canonical sequences, Aβ and tau as well as other amyloid forming sequences, like α-synuclein,TDP43, and TMEM106b. Amyloid fibrils are intermolecularly hydrogen bonded β-sheet protein aggregates containing parallel, in-register strands and are also characteristic features of many other neurodegenerative diseases (2). Monoclonal antibodies against Aβ amyloid are the first disease-modifying treatments to be approved for the treatment of AD. Aducanumab (Aduhelm), Lecanemab (Lequembi) and Donanemab (Kinsula) have been approved by the FDA as therapeutics to slow the progression of the disease with reported therapeutic activity of slowing clinical and functional decline by between 22-35% depending on tau burden (3). Although the approval of these antibodies validates Aβ as a therapeutic target there is considerable room for improvement in terms of effectiveness, dosage, side effects and cost (4).

Naturally occurring antibodies (NAbs) are believed to form the “front line” of protection against pathogens and infectious agents and are characterized by “polyreactivity”, or the ability to bind to many different protein sequences and “autoreactivity” or the ability to bind to endogenous protein sequences (reviewed in: (5-7). NAbs that react with selfmolecules (natural autoantibodies, NAuAbs) and pathogens occur in both healthy and diseased individuals (7). Aducanumab was derived from the blood of an elderly, cognitive normal human, indicating that disease modifying NAuAbs occur naturally in human blood (8). Additionally, there have been many investigations of human NAuAbs against Aβ as biomarkers and potential therapeutic agents (9). Unfortunately, no consistent conclusions have emerged from this work. There are many possible reasons for this inconsistency, including the fact that different antibodies may bind to different Aβ epitopes and it is not easy to distinguish among the many different antibodies that have different plaque binding properties that bind to the same amino terminal region of Aβ (10).

We have developed “Epitomic Profiling” (EP) as a tool for identifying the set of peptide sequences corresponding to epitope segments (ES) for natural antibodies (NAbs) in human serum and quantifying them (11-13). Epitomics is the analysis of the linear and discontinuous peptide ES that bind to antibodies that serves to define the epitope recognized by the antibody and as a fingerprint to distinguish binding mechanisms among different antibodies that bind to the same sequence, similar to epitope mapping by saturation mutagenesis (14). EP involves the immunoselection of 12mer random amino acid sequences followed by amplification of bound phage and panning two additional cycles followed by deep sequencing. The frequencies of antibody-specific ES are then compared in the AD and age matched, cognitively normal (NC) populations by Mann-Whitney U test followed by false discovery rate (FDR) correction to identify differentially expressed epitope segments (DEES). We recently applied EP to determine the set of tetrameric peptide ES that are associated with antibodies in serum samples from patients with the skin-blistering autoimmune disease, pemphigus vulgaris and healthy controls, which showed that different NAuAbs that target disease-associated proteins desmoglein 1, desmoglein 3 and the M3 muscarinic acetylcholine receptor are elevated in both disease and healthy control populations (13).

Here we report that human serum from both AD and cognitively normal controls (NC) have elevated levels of specific NAbs as determined by the number of DEES immunoselected. DEES recognized by NAbs in the NC population are 26-fold more abundant (FDR corrected p < 0.05) than DEES in the AD population, indicating that AD-specific Nab diversity declines dramatically with AD. Some of these DEES are contained within Aβ and tau sequences and the sequences of other amyloid-forming protein epitopes including α-synuclein, TDP43, TMEM106b and may be critical for preventing or delaying AD and may serve as prognostic or diagnostic biomarkers or potential therapeutic agents for AD. We also found that many of the DEES are highly correlated with each other corresponding to many different groups of antibodies that average approximately 50 DEES per group, consistent with the known polyreactivity of NAbs. If these groups correspond to a single polyreactive antibody or coordinately expressed antibodies, this suggests that many fewer polyreactive antibodies would be required to recognize all of the AD-associated DEES, compared to the number of monospecific antibodies produced by the adaptive humoral immune response.

## Material and Methods

We performed EP on 163 de-identified human serum samples obtained under an IRB-approved protocol from the UCI ADRC tissue repository for Alzheimer’s disease. Samples from 79 individuals were assessed by a panel of neurologists as AD and had Mini-Mental Status Exam, (MMSE) scores ranging from 1 to 30 mean 18.04±7.7,) and 84 samples from individuals over 50 years of age that were assessed as cognitively normal (NC) at the time of sample collection with MMSE scores that range from 27 to 30, mean 29.33±0.87), at the time of sample collection (SI_Table I. The EP approach relies on the immunoselection of random 12mers from a phage library, followed by elution of phage, amplification and then two additional panning steps as detailed in reference (13). DNA was isolated from the eluted phage, PCR amplified, bar coded and deep sequenced in a single run using Illumina NovaSeq sequencing resulting in approximately 3 billion total sequence reads distributed among 384 bar coded samples. The DNA sequencing files were then processed to extract and translate the random 12mer peptide sequences, identify unique sequences and count the number of times each sequence was observed and generate a two-column table of random sequences sorted by their observed frequency for each serum sample (Supplemental Information, SI_File1_Raw_12mers). The library is also sampled in triplicate in the absence of serum and sequences that bind specifically to protein A coated magnetic beads alone are filtered out of the 12mer sequences.

The 12mer sequences in SI_File1_Raw_12mers are then further processed by tiling them into 9 overlapping tetrameric sequences using a software routine written in Python. The purpose of tiling the random sequences is to consolidate antibody-specific sequences derived from different 12mer sequences and deconvolute the antibody-specific sequences from non-binding random sequences. We found that initial tiling in overlapping tetrameric ES is the optimal combination of sensitivity and specificity for the initial identification of ES that are recognized by antibodies that are differentially expressed in the AD and NC populations. We explored tiling in 3 residue epitope segments and while this approach is sensitive for finding antibody-specific ES trimers, it lacks specificity for initial tiling because only 8,000 different trimers are possible. This tiling approach is similar to a previously published “K-tope” approach that tiles the immunoselected sequences by pentamers (15), but tiling in pentamers lacks sensitivity and offers no advantage as larger linear segments, (e.g. pentamers, hexamers, etc.) are readily apparent as two or more sequential overlapping tetramers, so no specificity or uniqueness is lost by initially tiling in tetramers. Once an ES contained within a protein of interest has been identified, the presence of these longer ES can be readily verified by inspection of the sequence data in SI_File1_Raw_12mers. We also found that 4 residues is the most commonly observed as the size of an ES for anti-amyloid monoclonal antibodies that may be very different than antibodies against globular antigens (12).

After tiling the non-specific tetramers, which are defined as segments with a frequency that is indistinguishable from their frequency in the unselected library, are removed using Chi-square comparison of the observed and expected frequencies using a degree of freedom of 1 and p < 0.05. We tested the filtering algorithm by tiling random sequences from the unselected library and filtering them. We found that filtering removes an average of 80.6% of the tetramers derived from the non-selected sequences. The remaining unfiltered random sequences appear to be derived mostly from sequences that are over-represented in the library, perhaps due to artifacts of library construction or preferential amplification of phage bearing these sequences. The frequency of non-specific segments is set to zero in the output file (SI_File2_Counts).

The total sequence reads vary among individuals and is significantly lower in the AD group compared to NC (Supplemental Fig 1B). We examined whether this is correlated with IgG concentration in the samples, but ELISA measurements of IgG concentrations indicated that the IgG concentration in the AD group is not significantly different than the NC group (Supplemental Fig. 1C). A reasonable explanation for the difference in total sequence reads is the likelihood that the amount of cognate random sequence-bearing phage is insufficient to saturate the available antibodies in the AD group even after two subsequent amplifications as we observed that the diversity of NAbs in the AD population is much lower than that of the NC group, while the total IgG concentration is not significantly different between the groups. It is not clear what the best approach is to deal with this difference, so we examined the data with (SI_File3_Normalized) and without (SI_File2_Counts) normalization of the frequencies as the fraction of the total sequences. The two data sets give very similar results, but the normalized data set results in a fewer number of statistically significant ES differences between the NC and AD populations, so we focused on the data set normalized to the fraction of the total counts for subsequent analyses as a more conservative approach to minimize the differences unless otherwise noted in the text.

To evaluate the relationship between the frequency of an antibody binding ES and antibody concentration, we added increasing amounts of two different monoclonal antibodies, mOC1 and 4G8, that have well defined linear tetrameric epitopes DAEF (12) and VFFA (11) to human serum and determined the number of sequences corresponding to the specific epitope pattern we had previously determined for these antibodies. We found a linear relationship between the log fraction of sequences containing the cognate tetramer and the log antibody concentration, which corresponds to a power-law relationship between the fraction of sequences containing the cognate tetramer and antibody concentration. (SI Fig. 1A). Although the concentration dependence observed for the two antibodies is different, there is a range in which the number of antibody-specific sequences is linearly related to the antibody concentration. Therefore, the sequence frequency can be used as a surrogate for relative antibody concentration.

### Statistical analysis

Statistical analyses were performed using custom software coded in Python, in RStudio or in GraphPad Prism 10. We measured the distribution normality of the number of tetramers found on every individual sample using the D’Agostino & Pearson, Anderson-Darling and Kolmogorov-Smirnov tests in GraphPad. All three returned a non-normal result (SI_Figure 1 D, E), therefore we used the nonparametric Mann-Whitney U-test followed by multiple comparison false discovery rate (FDR) correction (16) to evaluate the tetramer frequencies that are significantly different between the AD and NC groups. The resulting FDR-corrected p values (q values) and mean rank differences among groups were used to generate volcano plots.

### WGCNA and UMAP analysis

In the Weighted Gene Co-expression Network Analysis (WGCNA) framework, the pairwise Spearman correlations between DEES are transformed into continuous edge weights by raising them to a power β. This transformation emphasizes stronger associations while attenuating weaker ones, producing a sparse yet information-rich network that approximates scale-free topology (17). The resulting network was visualized using a spring layout algorithm, in which spatial proximity reflects similarity in binding profiles. Node color indicates cluster membership as determined by hierarchical clustering, while node size scales with degree centrality to highlight potential hub peptides. Edge transparency and width are proportional to the underlying correlation strengths, visually emphasizing the importance of strong associations within and across peptide types.

Dimensionality was reduced by Uniform Manifold Approximation and Projection (UMAP) allowing complex relationships among samples to be visualized in a two dimensional space (18). UMAP was implemented via the ‘umap’ package in R, with default parameters and a fixed random seed to ensure reproducibility. The resulting two-dimensional UMAP embeddings were used as input for k-means clustering to identify subgroups within the data. The optimal number of clusters was determined using the average silhouette width method, evaluated across a range of 2 to 10 clusters (‘cluster’ package in R), with the silhouette width serving as a measure of clustering quality and cluster cohesion.

## RESULTS

### DEES are dramatically reduced in the AD population

EP identifies discrete linear and discontinuous peptide ES for antibodies that vary in concentration in human serum. We recently validated the ability of EP to identify and predict the binding sites for hundreds of NAuAbs associated with the autoimmune disease pemphigus vulgaris (13). We found that binding sites for disease-associated tetrameric ES covered over half of the protein sequence of desmogleins 1 and 3 and the M3 muscarinic acetylcholine receptor, which are known targets of disease NAuAbs. We also found that antibodies that are predicted to bind to the acetylcholine-binding loop of the M3AR receptor are able to activate the phospholipase C activity of the receptor in contrast to antibodies predicted to bind elsewhere. Here we used EP to identify Nab specificities associated with AD. We immunoselected random 12mer sequences from 79 AD and 84 NC serum samples and deep sequenced them as described in Materials and Methods. Briefly, unique 12mer sequences were identified and counted and output as a two-column format for each sample that contains the unique 12 amino acid sequence and the number of times it was observed (SI_File1_Raw_12mers). These sequences were then tiled into overlapping liner tetramers using the EP tiling module and the frequency of each tetramer in each sample counted and written to a file containing the observed frequency of each of the tetramers for each AD and NC serum sample (SI_File2_Counts). The purpose of tiling the random sequences is to consolidate antibody-specific sequences derived from different 12mer sequences and deconvolute the specific sequences from non-binding random sequences. The rationale for choosing initial tiling in tetramers is explained in materials and methods, but briefly it is the optimum combination of sensitivity and specificity for finding initial hot spots for antibody binding. Tiling in tetramers is also supported by a recent analysis of the geometric and physicochemical features of 1,309 atomic structures for antibody-antigen complexes in the SabDAb database. This analysis indicates that protein epitopes consisting of a single long linear segment are rare and that the vast majority epitopes are “conformational”, meaning that they consist of multiple discontinuous peptide ES that converge as a contiguous patch upon protein folding (19). The average epitope contains 14.6 ± 4.9 amino acid residues and 80% of epitopes contain between three and eight ES of 1-6 residues where longest ES has a minimum of 5 residues suggesting that we could expect to find at least one linear ES for most antibodies by initial tiling in tetramers (16). After tiling the 12mer sequences in tetramers and removing non-specific tetramers, the resulting data is written in a file containing the observed frequency of each of the 160,000 possible tetramers for each sample (SI_File2_Counts). This data was also normalized as a fraction of the total number of tetramers for each sample to correct for the lower number of sequences obtained from samples in the AD group compared to the NC group as described in Materials and Methods (SI_File3_Normalized). We chose the normalized tetramers data sets because they give more conservative results in terms of the differences between AD and NC DEES. We used the nonparametric Mann-Whitney U-test with multiple comparison FDR correction (16) to evaluate the ES frequencies that are significantly different between the AD and NC groups (Fig. 1A). A total of 18,940 DEES were found to be statistically significantly associated with AD (q < 0.05), with 696 DEES elevated in AD and 18,244 DEES elevated in NC after FDR correction (SI_File4 _DEES). The number of DEES observed in the NC population is much higher (approximately 26-fold) than in AD, indicating that diversity of disease-associated antibody specificities is dramatically reduced in AD. The overall antibody diversity as assessed by the number of unique, non-zero tetramers is lower in the AD population than in NC (mean value of 28,478 in AD vs 43,229 in NC, p < 0.007) (Fig. 1B), indicating that the 18,244 DEES elevated in the NC population can more than account for the decrease in total antibody diversity between the AD and NC populations.

**Figure 1:**
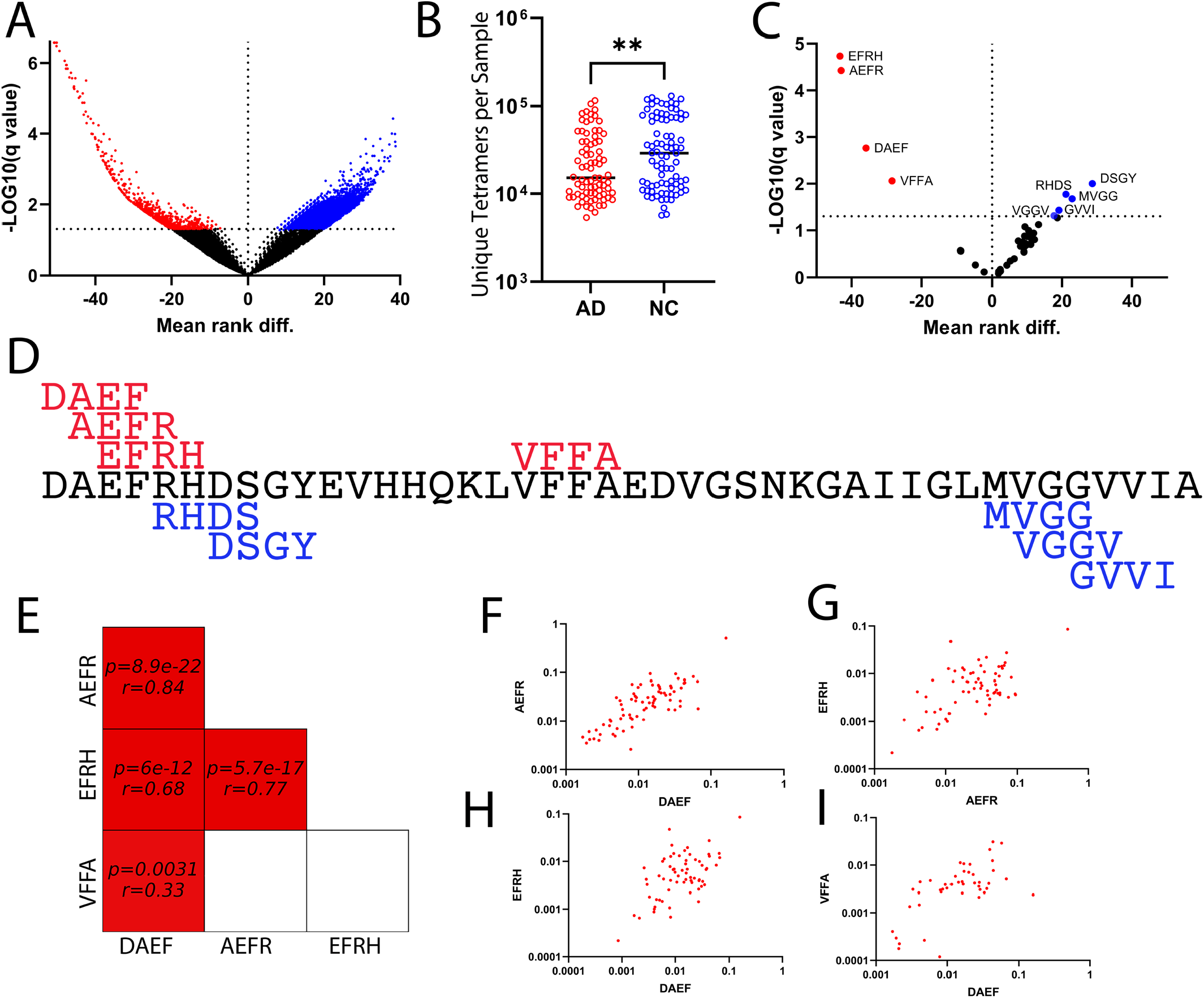
Disease-associated linear tetramer ES. A) Volcano plot of the DEES associated with AD. Red points are elevated in AD, while blue points are elevated in NC. B) Total unique tetramers in the AD (red) and NC (blue) sample populations ** p < 0.01. C) DEES that map to Aβ. The tetramers DAEF, AEFR, EFRH and VFFA are elevated in AD (red) while RHDS, DSGY, MVGG, VGGV and GVVI are elevated in NC (blue). D) Map of the locations of the DEES that map to Aβ. E) Pearson correlations for AD DEES that map to Aβ. F-I) Log-log scatter plots of the individual points for the Pearson correlations shown in panel E.

### DEES associated with amyloid sequences

After identifying DEES that are statistically significantly elevated in either the NC or AD groups, we analyzed DEES that map to the major amyloidogenic proteins observed in AD: Aβ, tau, TMEM106b, α-synuclein and TDP43 (20, 21) The DEES that map to Aβ amyloid are shown in Figure 1C, D. The Aβ DEES that are elevated in AD include 3 sequentially overlapping tetramers covering residues 1-6 (DAEFRH) and 18-21 (VFFA), while the DEES elevated in NC map to residues 5-10 (RHDSGY) and residues 35-41 (MVGGVVI) (Fig. 1D), indicating that different antibodies that target different Aβ epitopes are significantly elevated in both AD and NC. Some of the adjacent DEES overlap, suggesting that they may be segments of a larger linear epitope. We previously reported that EP is an effective and efficient approach for finding larger linear ES and discontinuous ES that is comparable to using tiled overlapping synthetic 10mer peptide arrays (12). We reasoned that we could exploit the natural 10,000-fold variation in observed antibody levels in populations to determine if the overlapping and discontinuous epitope segments are significantly correlated, which is evidence that they may be recognized by the same antibody. This would enable the building of larger, more complex epitopes that are more likely to identify a unique target by consolidating overlapping and discontinuous segments into a single epitope. If two or more tetramers are significantly correlated, then it is likely that they bind to the same antibody or to more than one antibody that are coordinately expressed. To test this, we conducted a Pearson correlation analysis of the DEES that map to Aβ and are elevated in the NC and AD populations (Fig. 1E). In AD, the overlapping DEES, DAEF, AEFR and EFRH are all cross correlated with Pearson r values ranging from 0.68 to 0.84 that are highly significant, indicating that they bind to an antibody that recognizes a hexameric linear epitope (SI_File5_amyloid_correlations). These predicted larger linear epitopes are easily validated by inspection of the 12mer sequence data (SI_File1_Raw_12mers), which shows that DAEFRH occurs in 39 sequence reads in the AD samples, while DAEFR and AEFRH occur independently much more frequently indicating that there may be multiple antibodies that bind this region. However, the discontinuous DEES VFFA is significantly correlated only with DAEF (r = 0.33; p < 0.0006), suggesting that these two DEES form a distinct conformational epitope. This suggests that there are at least two distinct groups of anti-Aβ antibodies in the population that are elevated in the AD population; one that binds to DAEF and VFFA and a group that binds to DAEFRH, DAEFR and AEFRH. Graphic representations of the pairwise ES Pearson correlation analyses indicate that the highly significant correlations consist of many non-zero points over a thousand-fold range (Fig. 1 F-I). The amino terminal residues 1-8 of Aβ are a hot spot for human anti-Aβ antibody reactivity (22) and we have previously found that 18 of 23 distinct monoclonal antibodies isolated from rabbits immunized with Aβ42 fibrils bind to at least one ES in this region (10). One antibody, mOC9, recognizes both DAEF and VFFA and several antibodies bind to DAEFRH (12), so antibodies with the predicted specificities in AD serum exist after immunization of humans and rabbits with Aβ42. In contrast to the AD DEES, none of the DEES that map to Aβ42 in the NC population shown in Fig. 1C are significantly correlated, including the adjacent DEES that overlap. However, we have described rabbit monoclonal antibodies, mOC76, mOC78 and mOC86, that have epitopes like those predicted for the DEES elevated in the NC population in Fig. 1D (12). Many DEES map to other amyloids found in AD brain, such as tau, α-synuclein and TDP43 (SI_File5_amyloid_correlations). Figure 2 shows the volcano plots of the FDR-corrected p values after comparing the tetramers that map to tau, α-synuclein and TDP43 (Figure 2A-C). Unlike Aβ, there are only three DEES elevated in the AD population for 2N4R tau, (SLAK, PGSE and TREP), two for TDP43, (QFOC, AEPK), one for a-synuclein (KTVE) and none for TMEM106B, even though these sequences are much longer than Aβ. For the AD DEES that map to tau, only two are highly corelated (PSGE at position 59 and SLAK at position 435, r = 0.645, p < 1.34x10^-10^) (SI_File5_amyloid_correlations). Unlike the NC DEES for Aβ that are all uncorrelated, most of the 48 NC DEES with q < 0.01 for 2N4R tau are significantly correlated, (SI_File5_amyloid_correlations). Only two pairs of DEES are highly correlated (KDQG, RHLS r = 0.82, and GSVQ, GGGQ r = .52,) while three DEES are all significantly correlated (DEGA, APVP, r = 0.45 and DEGA, KSPS, r = 0.4), consistent with the interpretation that they bind to three different antibodies that are typical epitopes consisting of 8 – 12 residues. However, 41 of the 48 NC DEES for tau are all cross correlated (r > 0.295) in multiple combinations (Fig. 2D, E). The DEES elevated in the NC population that map to α-synuclein and TDP43 show a similar grouping of a large number of correlated DEES (SI Figure 2Although the concentration dependence ). (SI_File5_amyloid_correlations), with α-synuclein containing a single cluster of 12 correlated DEES while TDP43 contains two clusters of 12 and 10 correlated DEES (r > 0.4). Amyloid fibrils like those from from Aβ, tau, α-synuclein, TDP43, and TMEM106b all contain a “fuzzy coat”, which is a region of sequence that is disordered and not part of the parallel, in-register β-sheet amyloid fibril core observed in cryoEM structures. If these highly correlated discontinuous DEES that map to natively disordered tau or its fuzzy coat belong to the same antibody, it suggests that these ES can coalesce together in close proximity to constitute a typical conformational epitope in many different alternative conformations of the disordered region.

**Figure 2:**
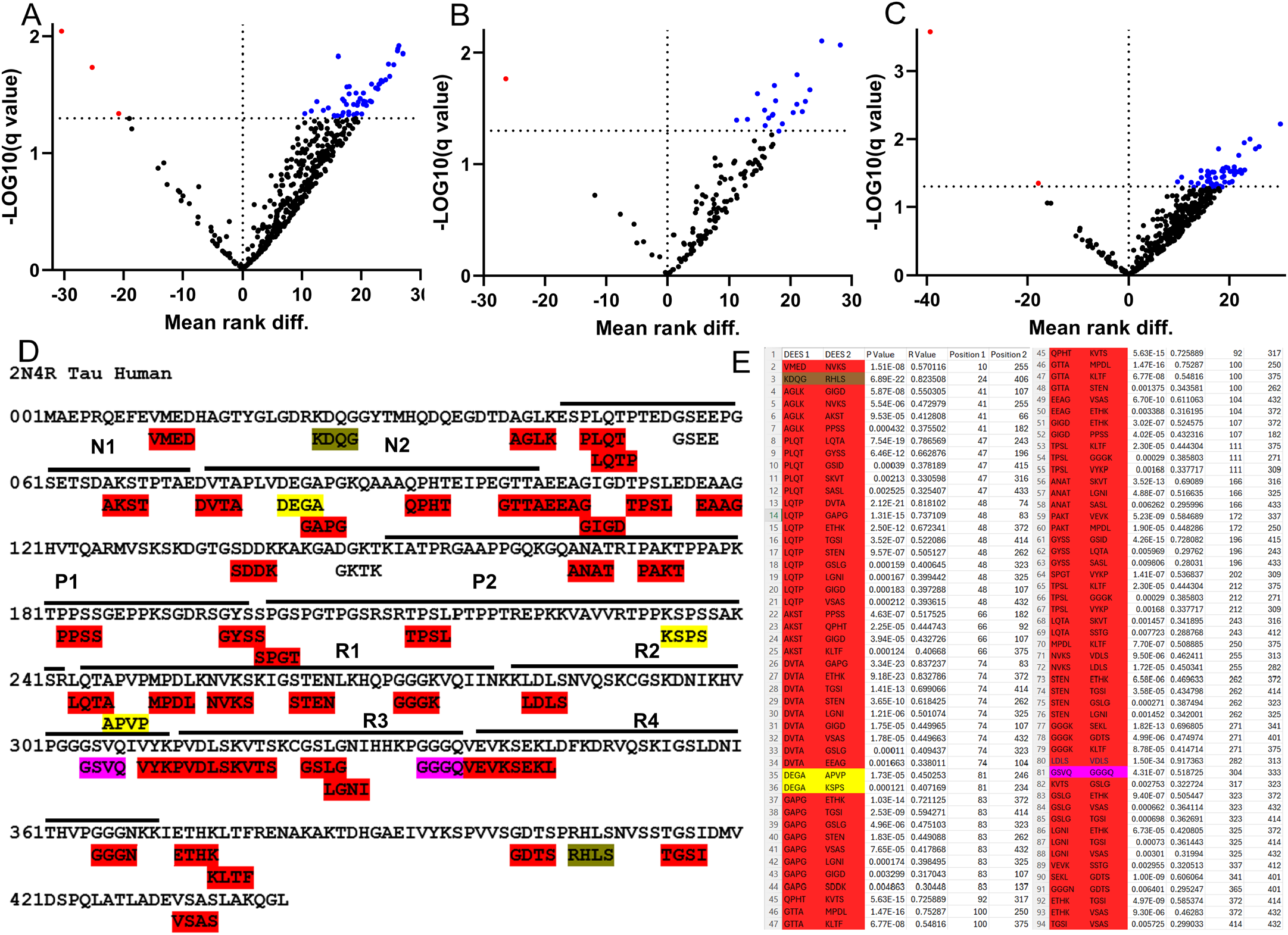
DEES that map to Tau, α synuclein and TDP43. A) DEES that map to 2N4R tau. B) DEES for α synuclein and C). DEES that map to TDP43. DEES that are elevated in AD are shown in red and elevated in NC are shown in blue. D) Location of the DEES that bind to 2N4R tau. The tau domains are indicated by lines over the sequence and labeled N1, N2, P1, P2, and R1-4. DEES that are highly correlated (p < 0.01, R > .295) are displayed in a colored font or highlighted in color. A group of 41 DEES that are all cross correlated are highlighted in red. E) List of p values and correlation coefficients (R) of the DEES shown in D.

### Highly correlated DEES in the AD and NC populations

Because so many NC DEES that map to tau are highly cross correlated, we examined how common this group co-correlation is in the wider sets of DEES for both AD and NC tetramers. We used all 696 of the significant AD DEES (q < 0.05), but to limit the number of comparisons, we only selected 784 NC DEES with q < 0.01 and we used both Pearson and Spearman approaches for calculating correlation coefficients. The two correlations tests give similar results. The highest 1000 correlations from both tests of the AD data set contain a total of 138 unique DEES and of this group 111 are common to both analyses. We chose to use the Pearson correlation data because the results are more conservative, yielding fewer statistically significant correlations with higher r values than the Spearman approach (3893 vs 8910, p < 0.01) (SI_File6_AD_DEES_Correlations). For the AD DEES, 154 pairs have an r value > .99 consisting of 62 unique tetramers in 7 co-correlated groups. This is an average of 22 correlated pairs per group and only two single correlated pairs exist. Groups 1 and 2 contain extensively overlapping sequence DEES, indicative of constituting longer linear epitopes as observed for Aβ (Fig. 1). Similar results were obtained for the DEES elevated in the NC population q < 0.01 although there is less extensive overlap indicative of long linear epitopes (SI Figure 3B, SI_File7_NC_DEES _Correlations). There are 261 pairs of NC DEES with r > .99 consisting of 124 unique DEES and 220 of the pairs fall into 14 co-correlated groups. In most groups there is no obvious sequence relationship and inspection of the 12mer sequences indicate that the two correlated DEES are not both discontinuous parts of a larger 12mer sequence. Group 3 contains the overlapping DEES: IQEP, QEPV, EPVQ, VQGR, GRTS, RTSK. This forms a large linear 11mer epitope IQEPVQGRTSK which is found in 54 different cells of the NC 12mer sequences (SI_File1_Raw_12mers). As we observed for the NC tau DEES, in most cases there are too many co-correlated AD and NC DEES to fit within a single antibody binding site, suggesting that either the DEES bind independently and alternatively to a single polyreactive antibody or they bind to a set of coordinately regulated antibodies in the population.

### WGCNA and UMAP Analysis of DEES in the AD and NC populations

To investigate the broader correlations between DEES in the AD and NC populations, we constructed a weighted correlation network encompassing both populations, where each node represents a DEES sequence and edges reflect pairwise similarity in binding profiles across msubjects. As in the Pearson correlation analysis above, we used all 696 significant AD DEES (q < 0.05) and 784 NC DEES where q < 0.01 (SI_File4 _DEES). Rather than applying a hard threshold like the Pearson correlations to define network edges, we adopted a soft-thresholding approach inspired by weighted gene co-expression network analysis (WGCNA) (17). By combining soft-thresholded network construction with hierarchical clustering, this framework provides a rich and unified view of the correlated DEES landscape, illuminating both shared and distinct patterns of immune reactivity in the AD and NC populations. This integrated network reveals a modular structure, including densely interconnected communities composed of both AD and NC DEES, as well as hub-like DEES that may serve as key immunological drivers or structural anchors (Fig. 3 A, D). Notably, several hub nodes bridged tetramer clusters, suggesting potential sequence continuity or functional complementarity. The WGCNA network of DEES in the AD population (SI_File8_AD_WGCNAnetworks), (Figure 3A-C) displays five distinct clusters containing the 696 significant DEES (q < 0.05). Cluster 5 forms the central hub of the network, characterized by dense intramodular connectivity and extensive links to other clusters. Many DEES in Cluster 5 show a high degree sequence overlap, correlation, centrality and strong interconnections, suggesting that they may coalesce as 3 larger linear epitopes that serve as key immunological associates of AD (Fig. 3B, C) along with partially overlapping DEES that reveal numerous wild card positions where substitutions of the canonical sequence are allowed (Fig. 3C). Epitope 1 of cluster 5 contains 13 amino acids and consists of the canonical sequence IGPLMHIMRHNPV. There are also 8 wild card positions at residues 3-9 and 12 in the sequence where some animo acid substitutions are allowed which are represented in ProSite notation as I-G-[PHI]-[LHI]-[MIRK]-[HQN]-[IN]-[MK]-[RQW]-H-N-[HPT]-V. A Blast Search using the canonical sequence against the clustered non-redundant protein data base returned several 9-10 amino acid hits with one mismatch. There is one 8 residue exact match, GPLMHIMR, which is a histidine sensor chemotaxis protein CheA from *Oceanidesulfovibrio marinus* (WP_167512608) and several other CheA chemotaxis proteins from other bacterial taxa. A ProSite Expasy scan using the wild card sequences [PHI]-[LHI]-[MIRK]-[HQN]-[IN]-[MIK]-[RQW] with multiple amino acid substitutions allowed in 7 positions returns the top ranked hit of PLIHIIR from (O25153 CheAY-HELPY) sensor histidine kinase from *Helicobacter pylori*, a human microbiome organism that has been implicated in increasing risk of AD (23). This protein is structurally and functionally related to the other bacterial CheA sequences identified in the blast search. Epitope 2 in cluster 5 is an 11 amino acid linear canonical sequence IGPHGEPKGGL that shares the first 3 residues with epitope 1 with two wild card positions at 5 and 8. A BLAST search of the clustered NR data base returns several 8-9 residue exact matches of predominantly bacterial origin. When restricted to human sequences, the top matches include several repeats within α-2 type XI collagen (AAA35498.1). Epitope 3 consists of DEES that map to the amyloid Aβ 1-6 region of the human amyloid precursor protein (APP), including wild card substitutions in positions 1, 2, 4 and 6, indicating that the AD DEES that map to Aβ are also part of cluster 5. Cluster 4 contains no overlapping DEES. Cluster 3 contains overlapping DEES that form the wild card heptamer sequences [HNK]-N-[WC]-F-P-[LI]-D. Cluster 2 contains the pentameric sequences ANFQL, HTTEI, and the wild card octamer M-I-[RL]-[DY]-H-P-F-T while Cluster 1 contains no overlapping DEES.

**Figure 3:**
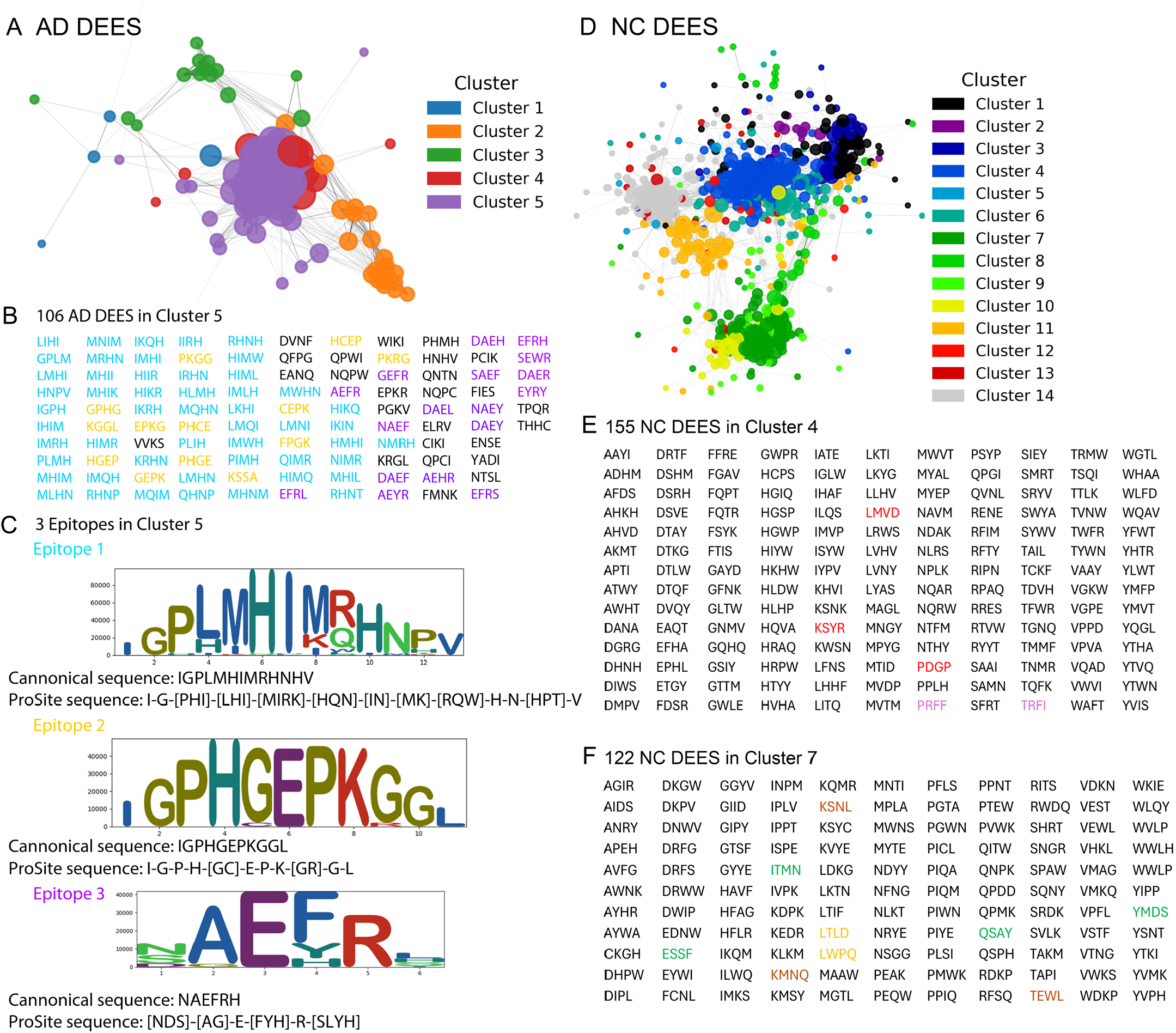
Weighted gene co-expression network analysis. A-C) WGCNA clusters of DEES elevated in AD. A) The 696 AD DEES (q < 0.05) form 5 co-expressed clusters. B) Cluster 5 contains 106 ES with the lowest q values that drive the association with AD. C) Most of the cluster 5 ES overlap to form 3 larger linear epitopes that each contain several wild card positions. The frequency of amino acids observed at each position is plotted as a LogoPlot for epitope 1 (blue), 2, (yellow) and 3 (purple). Epitope 3 maps to the amino terminus of Aβ. D-F) WCGNA clusters of NC DEES. The 784 NC DEES with q < 0.01 form 14 different clusters. D) Clusters 4 and E. cluster 7 have the highest number of the lowest q value DEES that are associated with normal cognition. Unlike cluster 5 in AD, there are few overlapping and highly correlated clusters that assemble into longer linear epitopes. The highly correlated (Rho > .99) DEES in clusters 4 (E) and 7 (F) are highlighted in colors to indicate their correlation.

The NC DEES fall into 14 clusters, including large clusters 4 and 7 (SI_File9_NC_WGCNAnetworks), (Fig. 3D-F). Unlike the cluster 5 for the AD DEES, there is less extensive sequence overlap of the tetramers that is indicative of forming longer linear epitopes. Cluster 7 contains 14 pentameric sequences and 1 hexameric sequence, while Cluster 4 has 8 pentamers and 1 hexamer. Some of the NC DEES that are highly correlated (SI_Fig3B, r > .99) are elements of clusters 4 and 7 as indicated by the same color code in Fig. 3E, F. The LogoPLots in SI_Files 8 and 9 show overlapping and wild card position substitutions with frequencies derived from the AD and NC populations (SI_File2_Counts).

### UMAP analysis of clusters in the AD and NC populations

To explore the distribution of DEES clusters within the AD and NC populations, we applied Uniform Manifold Approximation and Projection (UMAP) embedding of the DEES in AD, NC and combined AD and NC populations using the same DEES used in the WGCNA analysis. (Fig. 4). The AD DEES are contained within 3 clusters (SI_File10_AD_UMAP_Clusters) (Fig. 4A, D). Cluster 1 contains 18 subjects and is defined by the same DEES that make up epitopes 1-3 of cluster 5 of the WGCNA analysis. AD UMAP Cluster 2 contains 24 subjects and is driven by the partially overlapping DEES, AYMP MPSA and SAKI contained in WGCNA cluster 2. These tetramers overlap by 2 amino acids and appear to concatenate into a longer continuous amino acid sequence (AYMPSAKI), suggesting they may form part of the same extended epitope. However, this sequence does not occur in the raw 12mers (SI_File1_Raw_12mers). Rather, the 3 overlapping tetramers are part of a larger 12 amino acid canonical sequence, AYMPSASVSAKI containing a tetrameric insert ASVS that is by far the most abundant but contains many variants with one of more substitutions at residues 3-11. The predicted overlapping tile segments SASV, ASVS, SVSA AND VSAK are not found in the AD DEES because their q value is > 0.05. The sequence ASVS is the site of wild card positions with a large number of amino acid substitutions such that the set of tetrameric sequence variants in this region does not reach statistical significance. Thus, it is possible that these individuals in cluster 2 share a common antibody target or pathogenic target linked to this specific epitope structure. A BLASTP search of the canonical 12mer sequence returns no exact matches but several 8-11 amino acid hits that contain 2-3 mismatches. AD UMAP Cluster 3 is the largest with 37 samples and contains 11 DEES. It is driven by 3 DEES, ANFQ, NFQL which overlap to form a pentamer ANFQL and RDHP. This could represent a single discontinuous epitope, but ANFQ is not significantly correlated with either NFQL or RDHP, but NFQL and RDHP are highly correlated (r >0.99) (SI_File6_AD_DEES_Correlations).

**Figure 4:**
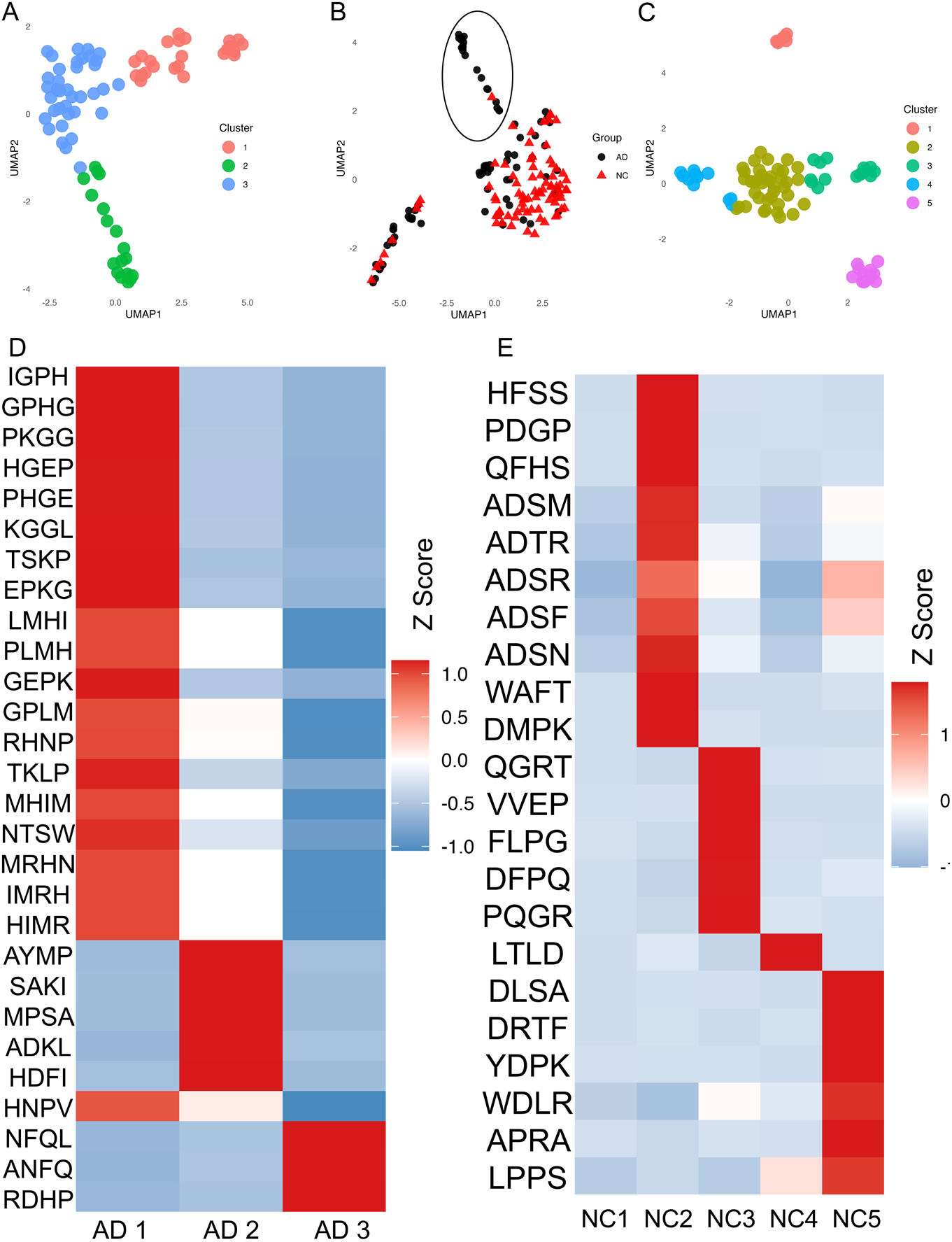
UMAP Analysis. A) UMAP embedding of AD DEES yields 3 clusters. B) UMAP Embedding of combined AD and NC DEES. The circled cluster consists mostly of AD individuals and is driven by the AD cluster 2 DEES AYMP, MPSA, and SAKI. C) UMAP clusters in NC. D) Heat map of the DEES that drive each AD cluster corresponding to panel A. Red indicates elevated levels and blue means reduced as shown by the scale bar. Each cluster is driven by a different set of DEES. E) Heat map of the DEES that drive each NC cluster corresponding to panel C. The largest NC cluster 1 has no specific DEES that are strong drivers of clustering.

When clustering was extended to include both AD and NC subjects, a smaller cluster (circled in Figure 4B) emerged that also showed elevated responses to AD cluster 2 DEES: AYMP, MPSA, and SAKI. This cluster included mostly AD subjects but also two NC individuals. The presence of this predominantly AD-like cluster that includes only two cognitively normal individuals suggests the two NC subjects may harbor latent or early-stage immune signatures resembling those in AD and suggests that epitope-based clustering may have the potential to identify subclinical immune phenotypes to aid early disease detection.

The UMAP embedding of the NC DEES (784 DEES q < 0.01) yields 5 clusters (SI_File11_NC_UMAP_Clusters). The DEES that drive each cluster are shown in the heat map (Fig. 4E). Cluster 1 contains 7 individuals, but none of the DEES in this cluster has as significant elevation level (>.0001). Cluster 2 is the largest containing 40 individuals and 63 DEES. Although the NC DEES show less extensive sequence overlap and long linear epitopes than the AD DEES, Cluster 2 contains DEES that overlap to form a heptamer of the ProSite sequence A-[AF]-D-[ST]-RMNK]-[FH]-S and a hexamer of the sequence YMVTMP. Cluster 3 contains 15 samples with 62 DEES that overlap to form 7 wild card sequences from 4 to 8 residues is driven by the hexamer sequence PQGR[TV]S and also includes the heptamer IQEPVQG and octamer SFYAMFQH. Cluster 4 contains 12 samples with 69 DEES that overlap for 9 distinct pentamers and is driven by the DEES LTLD and the wild card tetramer LPP[VS]. Cluster 5 contains 11 samples and 45 DEES including the pentamer A-D-[SH]-[RFM]-H. Several DEES are contained within more than one group, indicating that similar antibody specificities are distributed in more than one group. For example, the AD UMAP clusters 1 and 2, have GPLM, PLMH, LMHI, RHNP AND HNPV in common. For the NC UMAP clusters both Cluster 2 and cluster 5 contain the tetramer sequences ADSR, ADSF, ADSM forming A-D-S-[RFM] and clusters 4 and 5 contain LPPV and LPPS forming L-P-P-[VS]. The LogoPLots in SI_Files 10 and 11 show overlapping and wild card position substitutions with frequencies derived from the AD and NC population (SI_File2_Counts).

### Association of UMAP clusters with AD-associated clinical data

We also examined whether any UMAP cluster is associated with clinical or genetic parameters, like ApoE isoforms. Contingency table analyses were performed comparing cluster assignment with ApoE genotype and Mini-Mental State Examination (MMSE) categories using the Chi-square test of independence. No statistically significant global associations were observed between cluster membership and either ApoE genotype or MMSE score in either AD or NC cohort. However, examination of standardized residuals identified localized deviations from expected frequencies. Within the AD cohort, the ApoE2/3 genotype was modestly enriched in cluster 1 (standardized residual = 2.30), while in controls, the ApoE2/4 genotype demonstrated mild enrichment in cluster 3 (standardized residual = 2.63). These findings suggest the presence of subtle subgroup-specific trends that did not reach significance at the global association level and therefore should be examined in larger sample sets that contain more meta data.

## Discussion

We observed that there is a dramatic (26-fold) decline in the diversity of NAbs associated with Alzheimer’s disease. Although there is a generalized decline in immune function or increase in immunosenescence in aging (24, 25), the decline in diversity of antibody specificities is specific to AD because the samples examined here are approximately age matched populations (AD = 76.6 vs NC = 74.2 years). The overall diversity of antibody-specific ES in AD and NC is significantly different (43,229 ES in NC vs 28,478 in AD, p < 0.006) and the 18, 244 DEES elevated in NC can more than account for this difference.

NAbs are widely believed to represent the frontline defense against infectious agents and invasive pathogens and this is important for health, resilience and survival by neutralizing or clearing the infectious agent during the time it takes to mount an adaptive immune response (5, 26, 27). It is increasingly clear that the immune system plays a significant role in the development of AD and that monoclonal antibodies that target amyloid Aβ have disease-modifying activity in clearing amyloid deposits and slowing some measures of cognitive decline leading to FDA approval for AD (8, 26, 28). While some of the monoclonal antibodies have shown promising results, significant therapeutic benefit still seems to be illusive (4). The first FDA approved monoclonal, Aducanumab, was a Nab cloned from libraries of human B-cell clones derived from pooled blood from cognitively normal elderly subjects (8) and it recognizes residues 1-6 of Aβ. Our results suggest it would be much more likely to obtain such an antibody from AD subjects. The NAbs elevated in the AD and NC populations may play the same role in AD and AD related diseases as the monoclonal antibodies in human clinical trials. Several DEES map to the Aβ sequence within APP and distinct Aβ DEES are elevated in the AD and NC populations. The DEES elevated in AD cover residues 1-6 (DAEFRH) and 18-21 (VFFA), while the DEES elevated in NC map to residues 5-10 (RHDSGY) and residues 35-41 (MVGGVVI).

The antibodies that recognize DEES that are elevated in the AD population may arise from immune responses to disease processes, like amyloid deposition. The AD DEES map to the amino terminal residues 1-6 of Aβ, which is a hot spot for antibodies raised against fibrillar Aβ in both humans (22) and rabbits (12). The Aβ DEES elevated in NC overlap in sequence and map to residues 5-10 (RHDSGY) and residues 35-41 (MVGGVVI). Although the overlapping DEES would be expected to be derived from larger hexameric sequences, none of the Aβ NC DEES is highly correlated, suggesting that they may not bind to the same antibody. However a rabbit monoclonal antibody, mOC76, with a specificity corresponding to this predicted discontinuous epitope has been described (12).

Previous studies of NAbs in human serum or plasma that bind to Aβ have reported conflicting results with most of them using ELISA measurements and reporting that the levels of anti-Aβ antibodies are reduced in AD (reviewed in (29, 30) It is difficult to compare these prior studies to the results here as most of the studies have examined full length Aβ peptides or longer peptide segments within Aβ precluding comparison of specific epitopes, but our results are consistent with increases in different antibodies that recognize distinct epitopes in both the AD and NC populations.

DEES that map to other amyloids, like tau, TMEM106b and amyloids associated with subsets of AD, such as α-synuclein and TDP43 are also observed, but unlike Aβ, most of the DEES that map to these amyloids are elevated in NC, consistent with the dramatic decline in NAb diversity in AD. The tau NC DEES appear to demonstrate “polyreactivity”, a common property of NAbs, which is the observation that many of the DEES recognized are highly cross correlated as large groups or clusters implying that they may bind to the same antibody. Of the total 48 NC Tau DEES, 45 are cross correlated as a group with r > .29 (Figure 2, D,E). Of the 696 total AD DEES (q < 0.05) 62 are correlated into 7 groups with r > .99 and of the 784 NC DEES (q < 0.01) 123 DEES are correlated in 14 groups with r > .99. While some of these groups of correlated DEES show extensive sequence overlap because they are all derived from larger linear sequences, most of the DEES show no overlap. It is possible that the highly correlated DEES are discontinuous and non-overlapping but are still contained within the same 12mer sequence. However for the 100 correlation pairs with the highest correlation coefficients, none of the two DEES are contained within the same 12mer sequences.

The average number of amino acid residues that constitute an epitope is approximately 15 (19), so this would appear to be too many ES for them to coalesce simultaneously to all form the same epitope. If all of the highly correlated DEES bind to the same antibody, this could reflect the known polyreactive propensity of NAbs, but the individual DEES show little sequence or composition relationships that could account for them all binding to the same paratope. A study on the molecular basis for the polyreactivity of 6 monoclonal NAbs have shown that most bind to the same 6 distinct protein sequences with comparable, although relatively low affinities (31). They also found that no single amino acid residue, pattern or biochemical property like hydrophobicity or charge can account for the polyreactivity. However mutations that disrupt the binding to one antigen generally disrupt the binding of all tested antigens. Here we find a complementary scenario for the binding of correlated ES with no obvious amino acid composition or pattern to explain their co-correlation in large groups that may all bind to a polyreactive antibody. An EP analysis of polyreactive monoclonal Nab specificity might clarify the molecular basis for the polyreactivity.

An alternative explanation is that there are a set of NAbs in the NC population that are highly coordinately expressed. A study of the relationship between genetic variants in the IGH locus and the expressed IGH repertoire concluded that IGH germline variants determine the expressed antibody repertoire and that a single gene usage QTL variant can regulate the usage of multiple IGH genes (32). Both the individual polyreactivity of single antibodies and the coordinate expression of sets of antibodies may contribute to the disease-associated antibody repertoire and the response to disease pathogenesis.

The AD and NC DEES and the antibodies they bind to identified here may play different roles in disease and may be valuable for several different applications. Some may be useful biomarkers for disease onset and progression. The NC NAbs may normally recognize misfolded proteins and transmissible amyloid oligomers or prion-like seeds and bind and neutralize them to prevent amyloid nucleation and promote their clearance (33). In this mechanism, the NAbs associated with AD would have the same function ascribed to the NAbs that are proposed to represent the front line defense against more conventional infectious agents. It is possible that antibody binding may also promote the refolding of misfolded amyloidogenic sequences (34). The DEES may also be useful for screening for antibodies that bind to potential targets as therapeutic agents.

## Limitations

EP has broad potential for understanding the relationships between the NAb repertoire and health and disease, but it has a number of limitations. Since it is relatively new, it is also worthwhile considering approaches to overcoming or minimizing them. Since EP is based on screening random peptides, it is limited to epitopes that are peptide based, although it is possible it may find mimotopes of other classes of antigens. Currently no post translationally modified epitopes can be identified, but this may be possible for enzymatic modifications by splitting the library and modifying one population and then screening for DEES found in the modified library that are absent in the control unmodified library and checking if the amino acid at the position agrees with the enzymatic modification.

EP is based on the structural features of antibody-antigen binding contained in the SabDab data base, which indicates that 80% of the contact patterns contain at least one contiguous sequence segment of 3-8 residues, so more complex epitopes with more gaps and larger gaps would not be apparent in the approach described here. Epitope segments that contain few critical residues for binding may also not be apparent because they bind weakly or not at all. We have tested alternative tiling patterns containing 1 or more wild card positions of varying length as an approach to identifying more complex epitopes and this appears to be a fruitful alternative. Not only can this find more complex ES with variable gaps, it is also possible to find structure specific patterns associated with α helices and β strands such as hexamer with significant residues at positions 1, 2 and 5, 6 with intervening wild card positions at 3 and 4 that would be expected to bind to one side of an α helix that has 3.6 residues per turn and a β sheet heptamer with significant residues at 1, 3, 5 and 7 with intervening wild card positions at even residues. This pattern would be expected to bind to one side of a β strand.

EP is an antibody discovery platform that can identify a large number of DEES, so there is a problem with understanding the potential protein targets of these antibodies and choosing the ones of interest. The potential proteins can be identified in an unbiased approach by tiling the proteome of interest in the same tiling patterns as the random sequences and searching the tiled proteome with a selected list of DEES. The targets can be further triaged based on the known association of the predicted target with disease and by the complexity or uniqueness of the predicted epitope. EP can predict more complex epitopes assembled from overlapping tetramers and correlated discontinuous segments but need to be validated. Simple validation could include binding of the antibody of interest to synthetic peptides or recombinant proteins containing the epitope. More complex validation strategies can be based on the known function of the protein and testing whether NAbs that are predicted to a functional site have an effect on the protein activity as we have demonstrated for NAbs associated with the autoimmune disease pemphigus vulgaris where we show that samples with high levels of antibodies predicted to bind to the acetylcholine binding loop of the muscarinic M3 acetylcholine receptor activated the receptor to the same extent as acetylcholine while antibodies predicted to bind farther away had a reduced effect and samples that did not contain the predicted antibodies showed no activation (13).

In summary, our analyses provide a comprehensive landscape of the antibody repertoire diversity associated with normal cognition and AD and show that many of the ES associated with disease are either coordinately regulated at the level of sets of antibodies expressed or as a consequence of individual antibody polyreactivity, which implies that fewer antibodies may be necessary for maintaining normal cognition than would be expected if they act like typical monospecific antibodies. EP analysis may be applicable to many other diseases and health maintenance problems, like assessing the adequacy of the immune response to vaccination and predicting the susceptibility to infectious agents.

## Acknowledgements

We thank the UC Irvine Alzheimer’s Disease Research Center tissue repository for providing the di-identified serum samples. This work was supported by a grant from the Cure Alzheimer’s Fund to C.G.G.

## Author contributions

**Charles Glabe:** Conceptualization, Formal Analysis, Funding Acquisition, Project Administration, Methodology, Visualization and Writing the Original draft

**Joshua Netanel:** Formal Analysis, Writing – Review and Editing

**Jorge Mauricio Reyes-Ruiz:** Data Curation, Formal Analysis, Investigation, Methodology, Validation, Visualization, Writing -Original Draft

**Zhaoxia Yu:** Conceptualization, Formal Analysis, Methodology, Supervision, Writing – Original Draft

**Jizhi Zhang:** Formal Analysis, Visualization, Writing – Original Draft

